# Trade-offs between immunity and competitive ability in fighting ant males

**DOI:** 10.1101/2023.01.30.526206

**Authors:** Sina Metzler, Jessica Kirchner, Anna V Grasse, Sylvia Cremer

**Affiliations:** ISTA (Institute of Science and Technology Austria), Am Campus 1, 3400 Klosterneuburg, Austria

**Keywords:** male-male competition, male fighting, immunity-reproduction trade-off, immune gene expression, social immunity, inclusive sexual selection, social insects, *Cardiocondyla* ants, *Metarhizium* fungus

## Abstract

**Background:** Fighting disease while fighting rivals exposes males to constraints and tradeoffs during male-male competition. We here tested how both the stage and intensity of infection with the fungal pathogen *Metarhizium robertsii* interfered with fighting success in *Cardiocondyla obscurior* ant males. Males of this species have evolved long lifespans during which they can gain many matings with the young queens of the colony, if successful in male-male competition. Since male fights occur inside the colony, the outcome of male-male competition can further be biased by interference of the colony’s worker force.

**Results:** We found that severe, but not yet mild, infection strongly impaired male fighting success. In late-stage infection, this could be attributed to worker aggression directed towards the infected rather than the healthy male and an already very high male morbidity even in the absence of fighting. Shortly after pathogen exposure, however, male mortality was particularly increased during combat. Since these males mounted a strong immune response, their reduced fighting success suggests a trade-off between immune investment and competitive ability already early in the infection. Even if the males themselves showed no difference in the number of attacks they raised against their healthy rivals across infection stages and levels, severely infected males were thus losing in male-male competition from an early stage of infection on.

**Conclusions:** Males of the ant *C. obscurior* have evolved high immune investment, triggering an effective immune response very fast after fungal exposure. This allows them to cope with mild pathogen exposures without cost to their success in male-male competition, and hence to gain multiple mating opportunities with the emerging virgin queens of the colony. Under severe infection, however, they are weak fighters and rarely survive a combat already at early infection when raising an immune response, as well as at progressed infection, when they are morbid and preferentially targeted by worker aggression. Workers thereby remove males that pose a future disease threat by biasing male-male competition. Our study thus revealed a novel social immunity mechanism how social insect workers protect the colony against disease risk.

## Background

Escalated male fighting over access to females is common among many vertebrate species (Clutton-Brock *et al*., 1979), and can also occur in invertebrates, like cephalopods (Schnell *et al*., 2015), butterflies (Kemp and Wiklund, 2001), parasitoid and fig wasps (West *et al*., 2001; Abe, Kamimura and Shimada, 2005), as well as some ant species (Kinomura and Yamauchi, 1987; Heinze and Hölldobler, 1993; Heinze, Hölldobler and Yamauchi, 1998). Despite the high costs that fights can bear for both competitors, even lethal combats evolve when it allows the winning males to monopolize mating with the females (Clutton-Brock *et al*., 1979).

Male fighting ability relies on physical strength, which often becomes compromised when males get sick. Mounting an immune response to fight infection requires resources that cannot simultaneously be invested into fighting rivals, leading to a classic trade-off situation. Under infection, reallocation of resources away from growth and reproduction are therefore commonly observed already at early stages of infection like the incubation period before typical disease symptoms appear (Sheldon and Verhulst, 1996; Schwenke, Lazzaro and Wolfner, 2016; Barribeau and Otti, 2020). When disease has progressed, hosts often even show extensive “sickness behaviours” like lethargy and disengagement from most social interactions (Alexander and Stimson, 1988; Zuk *et al*., 1990; Hillgarth and Wingfield, 1997; Mazur and Booth, 1998; Eisenegger, Haushofer and Fehr, 2011), thereby conserving energy to fight off the infection (Hart, 1988; Kent *et al*., 1992; Dantzer, 2001; Shakhar and Shakhar, 2015). However, refraining oneself from engaging in male-male competition when sick does not seem a good strategy for males that will not gain a later chance for reproduction. This is the case either when disease will end deadly, or when hosts are so short-lived that reproduction cannot be postponed to after they have overcome the disease (Liu *et al*., 2017). Under such conditions, males have nothing to lose and often even fight more violently against a rival, as a terminal investment strategy (Clutton-Brock, 1984). Due to these apparent trade-offs, short-lived species typically do not invest many resources into a functional immune system, and in longer-lived species, immunity is often reduced during the reproductive period (also due to a direct immunosuppressive effect of the male hormone testosterone) (Alexander and Stimson, 1988; Zuk *et al*., 1990; Hillgarth and Wingfield, 1997; Mazur and Booth, 1998; Eisenegger, Haushofer and Fehr, 2011). Moreover, increased irritability and aggression are common behavioural reactions to disease in vertebrates, including humans (Hart, 1988; Loehle, 1995), which can also be found in insects, including ants (Konrad *et al*., 2018). Therefore, it is not straightforward to predict the effect of infection on male fighting ability, particularly since it will also be affected by (i) the level of infection, i.e. if it represents a low-level infection that causes mild or no disease symptoms, or a high-level infection inducing severe disease and death, and (ii) on its stage after exposure, i.e. whether the male suffers from progressed infection or is still in the early incubation period mounting up its immune response.

When fighting occurs in a social context, it may be further influenced by conspecifics. For example, the workers of social insects would have the potential to bias the outcome of male-male competition, if it takes place inside the colony. Such intra-nest fighting has evolved in several ant species, where mating occurs in the maternal nest and where emerging queen numbers are small enough to be monopolized by the winner male (Kinomura and Yamauchi, 1987; Heinze and Hölldobler, 1993; Heinze, Hölldobler and Yamauchi, 1998; Heinze *et al*., 2006). Whilst it is not yet known if such worker manipulation of male fights exists, it was found that ant workers can affect sexual selection by choosing or preferring some males as the queen’s partners (Sunamura *et al*., 2011; Helft, Monnin and Doums, 2015; Vidal *et al*., 2021). As workers are sterile but gain inclusive fitness via the reproduction of their related queens and males, such interference with mate choice has been referred to as “inclusive sexual selection” (Helft, Monnin and Doums, 2015). Biasing the outcome of male-male competition would similarly allow workers to favour some males over others as the queens’ future mates. In addition, worker aggression towards infected males could represent a form of social immunity (Cremer, Armitage and Schmid-Hempel, 2007). By helping the healthy male to win the fight, the diseased rival, who poses an infection risk to the colony, would be eliminated, reducing the danger of a disease outbreak in the colony. Such bias introduced by workers on male-male competition would therefore correspond to an ‘inclusive form’ of the Hamilton-Zuk hypothesis of parasite-mediated sexual selection (Hamilton and Zuk, 1982; Zuk *et al*., 1990).

We used the ant *Cardiocondyla obscurior* as a model system to test for the occurrence of worker interference with male-male competition in ants, and its possible interplay with male disease state. As typical for invasive ants (Holway *et al*., 2002; Cremer *et al*., 2008; Kenis *et al*., 2009), the queens and males of *C. obscurior* mate inside the nest (Heinze and Hölldobler, 1993; Heinze *et al*., 2006; Heinze, 2017). The wingless fighter males patrol the nest to identify emerging virgin queens to mate with, as well as emerging rival males to attack and engage in lethal combat. During a fight, males grab and besmear the rival male with a hindgut secretion that attracts workers, which then bite the besmeared male to death (Yamauchi and Kawase, 1992; Cremer *et al*., 2012). Besmearing is typically performed by the older and hence dominant male, yet fights between similarly-aged rivals often involve mutual grabbing and besmearing attacks (Cremer *et al*., 2012), which will give workers the opportunity to bias the fight outcome.

Successful males that are able to quickly detect emerging rivals thus can gain access to virgin queens over a long period of multiple weeks and can perform up to 50 and more matings (Metzler, Heinze and Schrempf, 2016). This has led to the evolution of both, a prolonged lifespan comparable to a worker’s lifespan, as well as a lifelong spermatogenesis in these fighter males (Heinze and Hölldobler, 1993; Schrempf *et al*., 2016). *C. obscurior* fighter males therefore greatly differ from ‘typical’ ant males, which are very short-lived, have completed their spermatogenesis before adult emergence and only have a single mating opportunity per lifetime in a mating flight, during which they engage in scramble competition with the other males (Hölldobler, B. and Wilson, 1990; Boomsma, Baer and Heinze, 2005; Boomsma, 2013). We would thus expect *C. obscurior* males to invest in a functioning immune system – in contrast to other social insect males with reported low immunocompetence (Gerloff, Ottmer and Schmid-Hempel, 2003; Vainio *et al*., 2004; Baer *et al*., 2005). Yet, this could make them vulnerable to an immunity-reproduction trade-off.

We therefore observed how exposure of *C. obscurior* males to the fungal pathogen *Metarhizium robertsii* affects the male’s own fighting ability and the workers’ interference in fights between a healthy and an infected male. To test for a possible trade-off between immunity and fighting ability we either exposed males to a low or a high dose of the fungal pathogen. Since the outcome of *Metarhizium* infections are dose-dependent (Hughes, Eilenberg and Boomsma, 2002; Konrad *et al*., 2012), the low dose was chosen to induce only a mild infection that the male’s immune system was expected to be able to cope with, whilst the high dose should in most cases cause severe disease, leading to death. For both, low and high dose, we tested the fighting success of infected males when in a combat situation with a healthy male (i) early in the infection, i.e. directly after exposure and when the pathogen’s infectious stages (the conidiospores, hereafter abbreviated as spores) penetrate the host’s cuticle, and (ii) at a progressed infection stage two days after exposure (Vestergaard *et al*., 1999). We expected that not only severe disease but already mounting an early immune response may negatively impact the males’ fighting ability. Apart from that, since the males face both a deadly disease and a deadly combat situation, they should show terminal investment of all their (remaining) resources into fighting. We also predicted that – if workers would indeed bias fight outcome – they should discriminate against the infected male, either to prevent pathogen spread by contagious males (early after exposure) or because the males represent a future infection risk to the colony by developing into a sporulating cadaver (when fighting at later stage of disease).

## Results

### Severe infections reduce male fighting success from an early disease stage

*C. obscurior* males that had been exposed to a high pathogen load of *M. robertsii* had a reduced fighting success, when in a 24h-long combat situation with a healthy male (exposed to a pathogen-free sham treatment; control male). This was equally so for the early stage of infection, when males were put into the fight situation immediately after pathogen exposure (Fig. 1a), and when their infection had already progressed to a later stage, as males had been exposed already 48h before start of the fight (Fig. 1b). Compared to fights between two healthy males, fights, in which one of the males was infected with a high dose, more frequently ended with one clear winner and one clear loser, i.e. with one rival surviving and the other being killed in combat (trending in early infection: GLMM χ^2^=4.258, df=1, p=0.059; late infection: χ^2^=4.936, df=1, p=0.039; Table S2; Figs. 1a,b inserts). Importantly, in the great majority of these fights, it was the infected male that lost the competition, independent of the stage of its infection: 85% (34/40) of the early-infected males (χ^2^-Test χ^2^=11.169, df=1, p<0.001), and 93.5% of the late-infected males (43/46; χ^2^-Test χ^2^= 21.445 df=1, p<0.001) had been killed at the end of the fighting period when exposed to a high pathogen dose (Fig. 1a,b).

**Figure 1.**
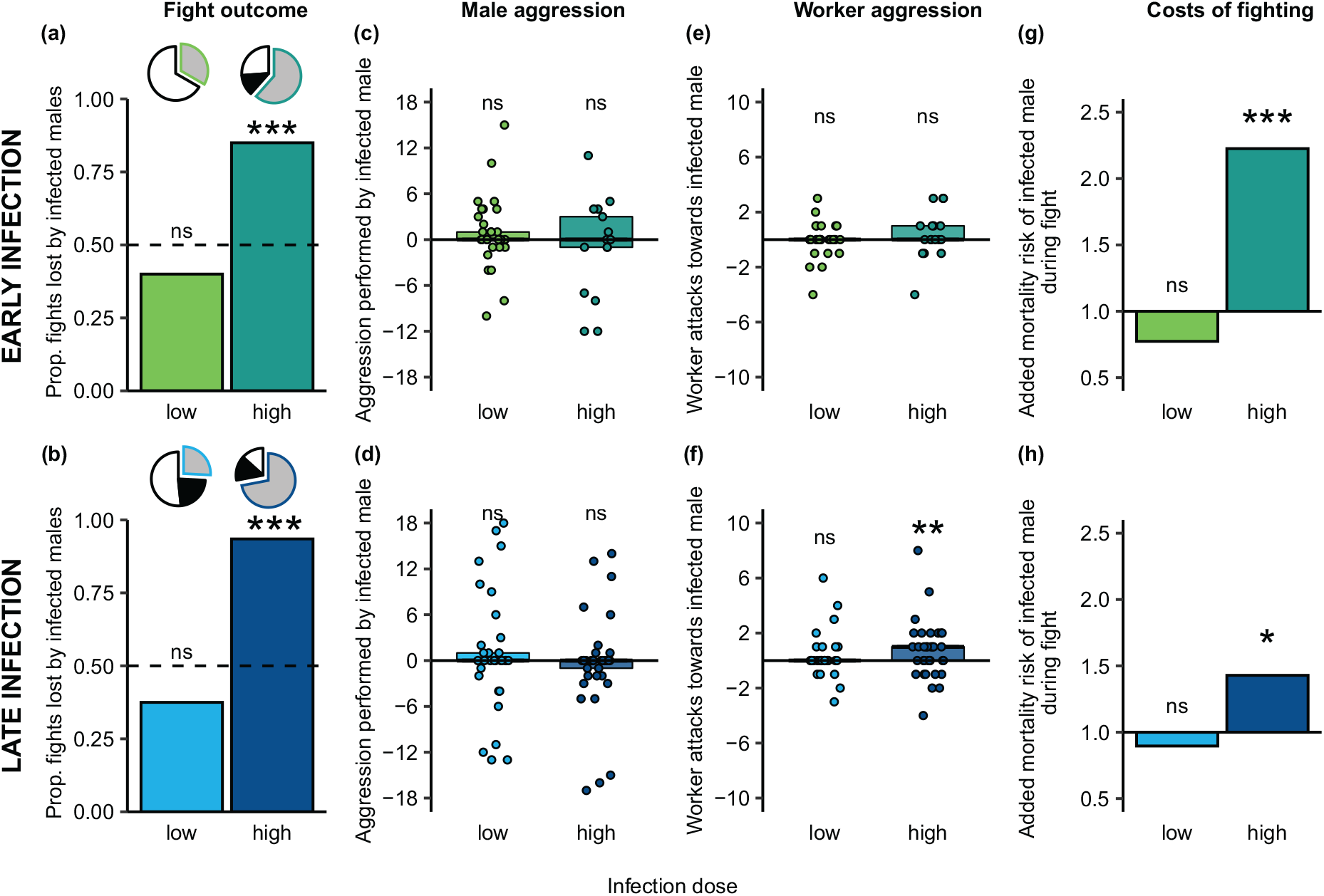
Fighting ability of infected males in dependence of infection dose and stage. Fight outcome for infected males compared to their healthy rival males at (**a**) the early (green) and (**b**) the late (blue) stage of infection. Inserts show the proportion of fights that ended undecided with either both males still alive (white), both males dead (black), or with one clear winner and loser (grey). In these decided fights (grey offset slice with coloured outline), infected males had a higher risk of losing the fight than by chance (dotted line) after exposure to a high, but not low pathogen dose (darker tone reflects higher dose; based on a total of 347 fights, Tables S1,S2). The aggression level of infected males (**c,d**) did not differ from that of their healthy rivals across infection stage or level, whilst workers showed increased aggression only to late-stage high-level infected males (**e,f;** Table S3). Individual data points depict aggression by (c,d) and towards (e,f) each infected male compared to its healthy rival (aggression_infected-healthy_, i.e. resulting in a zero value under equal aggression and positive values for higher aggression), boxes give the 95% CI and median as black line (based on 112 observed fights between healthy and infected males). The added risk of dying for infected males when fighting (**g,h**) compared to their baseline mortality when non-fighting during the same period (Fig. S1; Table S4) was significantly increased for males in the early and late stage of a high-level infection, where their mortality risk during fight increased 2.2-fold at the early stage and 1.4 fold at the late-stage (Table S5). Bars depict the fold change of mortality during combat to the males’ respective baseline mortality (value of 1 depicts equal mortality). Based on 201 fights. Significant deviation from chance (50:50) (a,b), from the healthy male (c-f) and from their non-fighting baseline mortality (g,h) given for each group. * p<0.05, *** p<0.001, ns=non-significant.

When males had only received a low pathogen dose that rarely causes disease, there was no difference between the probability of the fight ending with a clear winner and loser compared to fights between two healthy males, for both infection stages (early: GLMM χ^2^=0.189, df=1, p=0.236; late: χ^2^=1.125, df=1, p=0.433). Moreover, when fights were decided, low level-infected males did not lose the fights with a higher probability than their healthy rivals (early: Fisher Exact Test p=1.000; late: Fisher Exact Test p=1.000; Table S2; Figs. 1a,b).

### Workers attack severely-infected males at a late disease stage

To test whether the fighting success of infected males could be explained by their own aggression level towards their competitor, or by the aggression they received from the workers, we observed the behaviour of the fighting males and their workers. We found that – independent of dose and stage of infection – infected males showed the same level of aggression towards their rivals as the healthy males did towards them (Figs. 1c,d; GLMM all p>0.609; Table S3). This means that even the high-level infected males, who finally lost the fight in the majority of cases (Figs. 1a,b), did not perform fewer attacks. The number of worker attacks towards the infected male compared to its healthy rival was twice as high in fights with high-level exposed males at a late stage of their infection (GLMM χ^2^=0.1552, df=1, p=0.009), but not in the early stage, or after low-dose exposure of the males (Figs. 1e,f; Table S3).

### Males suffer highest fighting costs early in severe infection

Worker aggression could explain why high-level infected males at a late disease stage frequently lost the fights. However, at the early infection stage, the high-level infected males had equally low fighting success, even if workers did not treat them with increased aggression (Figs. 1a,b,e,f). We therefore determined the baseline mortality risk of the males in their respective fighting period in the absence of a fight, to disentangle, how much of the mortality of fighting males could be contributed to the fight itself, and how much to an already impaired survival due to infection. To this end, individual males were kept in the absence of a competitor with their workers for a 24h period, starting either directly after pathogen exposure, or after infection had been established for 48h. Whilst healthy males had a mortality risk of 8-9% (Table S4), infection in general increased this risk. At the early stage of infection, this was only significant for males exposed to a high pathogen dose (low dose: 17%; GLMM χ^2^=3.323, df=1, p=0.091; high dose: 29%; GLMM χ^2^=8.992, df=1, p=0.005). When infection had progressed, the males’ baseline mortality risk in the absence of fighting was already pronounced at the low (36%; GLMM χ^2^=7.624, df=1, p=0.010), and reached high morbidity (56% mortality) at the high pathogen dose (χ^2^=24.820, df=1, p<0.001; Fig. S1; Table S4). Notably, this means that progressed infection led to a high infection-induced baseline mortality, with no significantly further increase of mortality caused by fighting at the low dose (GLMM χ^2^=0.086, p=0.769), but at the high dose (1.4-fold increased mortality; GLMM χ^2^=7.107, p=0.010, Fig.1h, Table S5). Early-infected males showed comparably lower baseline mortality, but died more than two times more frequently during fighting when infected with a high dose (2.2-fold increased mortality; χ^2^=16.120, df=1, p<0.001; Fig. 1g, Table S5). Fighting costs therefore incurred for males in the early and late stage of severe infection, but were much more pronounced for early stage-infected males (Figs. 1g,h).

### Severe infection induces a strong early immune response

These data suggest that fighting as an additional stressor may unveil costs of mounting an immune response at the early stage of infection, causing increased mortality when the immune response is strong, as expected under severe but not yet mild infection. We therefore tested if immune genes were activated already within the first 24h after pathogen exposure, and whether this immune activation was higher after high-level exposure. Increased immune gene expression at the level of receptors and effectors would indicate that the infectious fungal spores were able to breach the host cuticle, cause infection and activate the host’s immunity cascade within 24h after exposure. We therefore measured the expression of three genes that act at different stages of the immune response, (i) Relish, a signal cascade gene in the IMD pathway (Myllymäki, Valanne and Rämet, 2014; Sheehan *et al*., 2018); (ii) Prophenoloxidase(PPO)-activating factor, which starts the process of pathogen melanisation (Gillespie, Kanost and Trenczek, 1997; Cerenius and Söderhäll, 2004; Cerenius, Lee and Söderhäll, 2008); and (iii) Defensin, an antimicrobial peptide destroying fungal pathogens (Viljakainen and Pamilo, 2005, 2008). We indeed found that all three genes were upregulated 24h, but – with the exception of Relish (Fig. S2) – not yet 12h after exposure to the high pathogen level (high dose 12h: Wilcoxon-Test, Relish p=0.045, all others n.s.; high dose 24h: all p-values < 0.027; Table S6; Figs. 2, S2). In the low-dose exposure, none of the genes was significantly upregulated compared to the baseline gene expression in healthy males (low dose 12h and 24h: Wilcoxon-Test, all p-values > 0.098; Table S6; Figs. 2, S2). These data are in line with the observed survival cost of mounting an immune response early after exposure, in particular when under the additional stress of combat.

**Figure 2.**
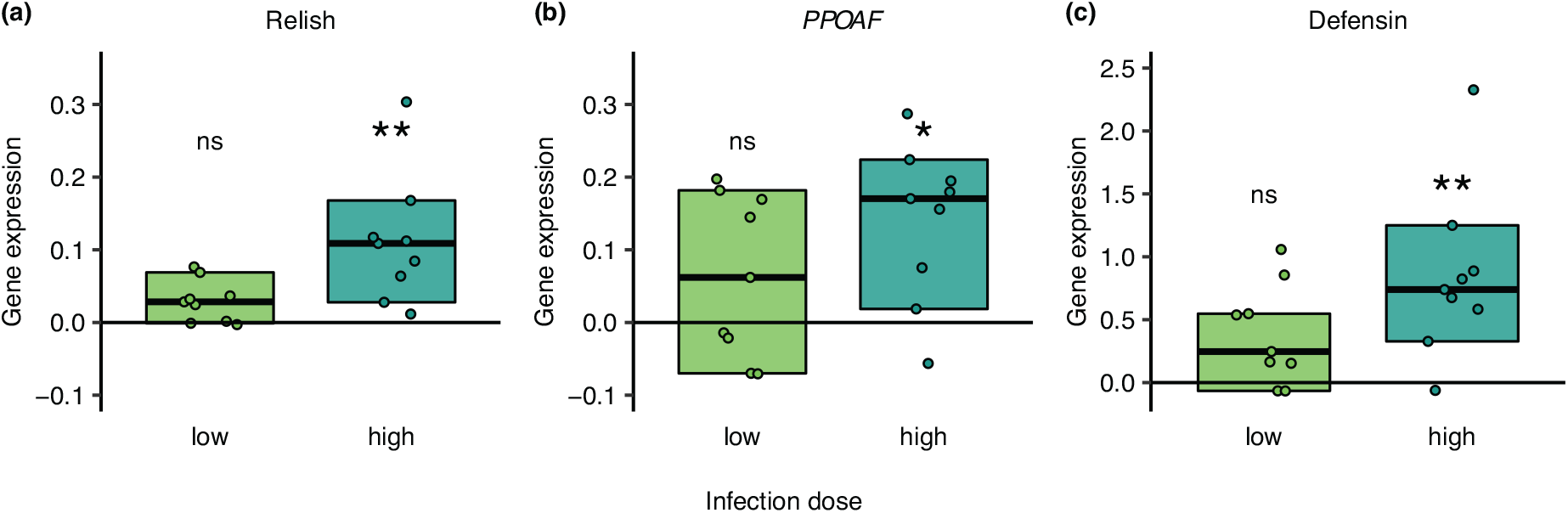
Immune gene expression 24h after exposure to the low or high pathogen dose. Expression of all three measured immune genes (**a**) Relish, (**b**) *PPOAF*, and (**c**) Defensin was increased in infected males compared to the baseline expression level in healthy males (zero baseline) for males after high-dose (dark green), but not the low-dose (light green) pathogen exposure. For each infected male we show the relative expression level of each gene normalised to the housekeeping gene *EF1*, relative to the median gene expression of the respective healthy control males (zero line) as individual data point, median per group as black line and 95% CI as box. Based on a total of 28 males (Table S6). Significant deviation to the healthy males for each dose and gene indicated by * p<0.05, ** p<0.01, ns=non-significant. See Fig. S2 for gene expression 12h after exposure.

## Discussion

We found that the competitive ability of fighting *C. obscurior* males was strongly impacted by severe (but not by mild) infection, and that these negative effects already occurred early upon exposure. This is in line with general observations across vertebrates and invertebrate species that both incipient and progressed infection can be costly (Sheldon and Verhulst, 1996; Schwenke, Lazzaro and Wolfner, 2016) for males and negatively affect their fighting ability.

High-level infected males were inferior to their healthy rivals and suffered a very risk of losing the combat and dying, both when entering the fight immediately after exposure to infectious *M. robertsii* spores, or with an already established infection. At this late infection stage, male morbidity was already very high, since >50% of the males would not have survived the day even without a fight. However, this was not yet the case during early infection, where male mortality risk more than doubled when confronted with a rival compared to when being kept without an opponent. This suggests a classic fighting-immunity trade-off (Schwenke, Lazzaro and Wolfner, 2016) during early infection, when *C. obscurior* males strongly invested into raising an immune response. Costs of this immune investment became aggravated when the males faced the additional stress of fighting against a rival. Mild infection, on the other hand, resulted in only a minor (and non-significant) immune gene upregulation, and males were not inferior in fights against healthy rivals. This is further evidence that resource depletion by mounting an immune response may cause the lower observed competitive success of males when fighting a severe infection.

Raising such a strong immune response is an unusual trait for males of social Hymenoptera, including the social bees, wasps and ants. These males typically have much lower immunocompetence than the colony’s workers or queens (Gerloff, Ottmer and Schmid-Hempel, 2003; Vainio *et al*., 2004; Baer *et al*., 2005), which is attributed to both genetic and life history effects. First, males of haplodiploid species are the haploid sex with lower variability on disease defence loci (O’Donnell and Beshers, 2004; Ruiz-Gonzáles and Brown, 2006), and, second, their lifespans are often only a few days, where even a good immune defence would not substantially increase life expectancy (Stürup, Baer and Boomsma, 2014). The high investment into immune defence of *C. obscurior* fighter males that we describe here likely evolved due to their uncommonly long lifespan of up to several weeks or even months (Metzler, Heinze and Schrempf, 2016), during which they can engage in multiple (up to approx. 50) matings with the emerging virgin queens (Metzler, Heinze and Schrempf, 2016). A functioning immune system may therefore be critical for males to ensure surviving at least mild pathogen threats at low cost to their fighting ability and giving them access to many more future mating opportunities. Therefore, we suggest that, even if a high-level infection (which will typically later develop into fatal disease) induces a trade-off with the male’s fighting ability already at an early infection stage, the benefit of having a functional immune system will outweigh these costs as it allows the males to overcome mild infections without making them inferior competitors. Multiple factors, such as (i) these males not leaving their maternal nest, (ii) *C. obscurior* ants nesting in arboreal cavities rather than in pathogen-rich soil (Heinze *et al*., 2006), and (iii) their active performance of hygiene and infection removal behaviours (Ugelvig *et al*., 2010), combine to make it likely that contamination of the males with a low pathogen level either via the nest or contagious colony members will be a more common scenario in natural colonies than infection of males with very high doses.

Interestingly, infection levels did not directly impair male aggression, as we could not detect any differences in the aggressive behaviours performed by infected males at any stage or intensity of infection compared to their healthy rivals. This indicates that even the moribund males at the late stage of a severe infection recruited their last resources into fighting their competitors, likely as a terminal investment strategy (Clutton-Brock, 1984). However, these males were more aggressed by workers than their healthy rivals, revealing that workers engaged differently into the fights depending on the health state of the competing males. This is in line with inclusive sexual selection (Sunamura *et al*., 2011; Helft, Monnin and Doums, 2015; Vidal *et al*., 2021), in which workers bias the outcome of male-male competition. A non-exclusive explanation could be that workers attack the fatally-infected males to eliminate them before they die and infectious spores grow out of their cadavers. Such “destructive disinfection” (Pull *et al*., 2018) represents an efficient social immunity measure in social insect colonies (Cremer, Armitage and Schmid-Hempel, 2007; Cremer, 2019), preventing the development of sporulating cadavers that would otherwise pose a very high risk for disease outbreak in the colony (Hughes, Eilenberg and Boomsma, 2002; Pull *et al*., 2018).

Workers did not discriminate against males directly after exposure, even if these males carried not yet firmly-attached infectious spores that we found to cross-contaminate the rival male in 45% (9/20) of the fights (N=20 fights of high-level exposed vs healthy males analysed for spore transfer). However, this was also not to be expected as aggression in social immunity rarely occurs at the very early stage of pathogen exposure, but rather when infection has established (Pull *et al*., 2018). Even though the males had very close contact during the fights, the contaminated rival carried on average less than 3% of the spore amount of the originally exposed male (500 vs 17,800 spores; see source data), which its immune system was likely able to cope with.

## Conclusions

Our study revealed that disease negatively impacts male competitive ability in fighting ant males. Interestingly, however, we found that the same outcome – losing the fight – had different causes during the early and late stages of infection. While late-stage males were already moribund at the time of combat and received increased aggression by workers, early-infected males invested heavily into mounting an immune response, which translated into a high mortality when they had to additionally fight a competitor male. Whilst this was true for severe infections, mild infections only led to a non-significant immune activation and did not induce any substantial fighting costs. This shows that the fighting–immunity trade-off is dose-dependent and only occurs at high infection levels, while a well-functioning immune system likely allows males to cope with low-level pathogen exposures in the nest without comprising their fighting ability and allowing future mating success. As being successful in fighting off rivals is key for the males to gain access to female mates, males invested all their energy into fighting, even at terminal disease stages. Under this – and only under this – condition, workers took side and over-proportionally attacked the already moribund male. This inclusive sexual selection by biasing the outcome of male-male competition represents a novel mechanism allowing workers to ensure colony health through social immunity.

## Methods

### Ant hosts

Colonies of the Myrmicine ant *Cardiocondyla obscurior* (Wheeler, 1929) were collected in 2013 and 2017 from Tenerife (28.390348 N, −16.596098 E; permits ABSCH-IRCC-ES-237603 ESNC2, Expte: AFF 199/17). Ants were kept in large stock colonies at constant 27°C and a day/night light cycle with 14h of light in the laboratory. Colonies were fed twice a week with minced cockroach and were supplied with water and 30% sucrose *ad libitum*. All experiments were performed at 27°C. Ant collection, rearing and experimental work was in line with European law and institutional guidelines.

This species shows a male diphenism of wingless fighter males that regularly occur in the nest and engage in deadly fights (Kinomura and Yamauchi, 1987; Heinze and Hölldobler, 1993; Heinze, Hölldobler and Yamauchi, 1998; Cremer *et al*., 2012), as well as peaceful winged disperser males that are produced under stress conditions (Cremer and Heinze, 2003). Since only the wingless males engage in fighting, we focused on these males in our study. Males were collected as pigmented ‘ready-to-hatch’ pupae from the brood pile and kept with two nurse workers (to help eclosion), until all males were between three and six days old, and their cuticle fully sclerotised when they entered the experiment. We used these age-controlled males to avoid differences between them in their *a priori* fighting ability, as otherwise old males have a clear fighting benefit over males younger than two days of age, which still possess a soft cuticle (Cremer *et al*., 2012).

### Fungal pathogen

As a pathogen, we used the fungus *Metarhizium robertsii* (strain ARSEF 2575) with an integrated red fluorescent (mRFP1) label (Fang, Pei and Bidochka, 2006), obtained from M. Bidochka, Brock University. *Metarhizium* fungi, incl. *M. robertsii*, are generalist insect pathogens that cause natural infections in ants (Pull, Hughes and Brown, 2013; Angelone and Bidochka, 2018; Casillas-Pérez *et al*., 2022). Their infectious stages, the conidiospores (abbreviated as “spores” throughout), attach to the body surface of their insect hosts, germinate and breach the host cuticle to cause internal infection shortly after. High infection doses cause host death within several days, followed by formation of new infectious particles from the sporulating cadaver (Hajek and St Leger, 1994; Vestergaard *et al*., 1999). Low-level infections rarely cause disease, and can even induce immune stimulation protecting against future infection (Konrad *et al*., 2012; Liu *et al*., 2015).

Prior to each experiment, spore suspensions were taken from long-term storage (−80°C) and cultivated on sabouraud dextrose agar (SDA) at 23°C for approximately three weeks. Conidiospores were harvested in 0.05% Triton X 100 (Sigma Aldrich; in autoclaved distilled water). Spore suspensions were counted with a Cellometer Auto M10 counter (Nexcelom) and adjusted to the respective concentrations needed for the experiment (see below). The germination rate of spore suspensions was confirmed to be >95% before the start of each experiment. To induce infection, males were shortly dipped into the spore suspension and placed on filter paper to absorb excess liquid. Healthy control males were set up in parallel to the fungal treatment in each experiment. They were treated the same way, except that they were only dipped into 0.05% Triton X (sham treatment).

### Experimental setup

To determine the effect of early infection, males were dipped into a suspension of either 10^6^ spores/ml (low dose) or 10^9^ spores/ml (high dose), and were placed with five workers and a second male from their colony, all sham-treated. For both the low and high dose, we equally set up control fights, where both males had only received the sham treatment (total 168 early-stage fights, detailed in Table S1). Workers immediately groom off spores from freshly-exposed colony members, so that the number of infectious spores that will then enter the body of the exposed host is drastically reduced (Rosengaus *et al*., 2000; Walker and Hughes, 2009; Reber *et al*., 2011). As spores on a freshly-exposed male are still removable, it is also possible that they cross-contaminate their nestmates (Konrad *et al*., 2012), including the rival male, during this period.

To obtain late infection stages, we therefore kept each male in isolation for 48h post exposure before placing them to the workers and rival male. During this period, the spores firmly attach and enter the body to cause internal infection (Vestergaard *et al*., 1999), so that males also no longer carry any infectious spores on their body (Walker and Hughes, 2009) when entering the fight. To prevent these males developing multi-fold higher infection loads due to the lack of spore removal by grooming workers, we used lower exposure doses for induction of these late-stage infections (low dose: 10^4^ spores/ml; high dose: 10^6^ spores/ml). Sham-treated control males were similarly isolated for 48h, and the workers kept for 48h in groups of 35 individuals until the start of the fight (total 179 late-stage fights, detailed in Table S1). All fights contained the two rival males and five workers, all originating from the same colony, and were performed in experimental containers (height 28mm, diameter 25mm) with a plastered and watered base to ascertain the required humidity. The fight outcome was determined 24h after putting the two males together and with their workers for all 347 fights.

For each combination of infection stage and level (early low and high, late low and high), and equally for the fights between one infected and one healthy male and for the control fights between two healthy males, we observed the fight outcome per fight (number of dead and alive males), as well as male and worker aggressive behaviour for a subset of fights (see Table S1 for sample sizes). In fights between a healthy and an infected male that had a clear winner (the surviving male) and a clear loser (the dead male; N=112 fights), we distinguished, whether it was the infected or the healthy male losing. To this end, we either quantified the spore load of both males at the end of the experiment (when males were not colour-coded; using droplet digital PCR, see below; N=48 fights, Table S1) – thereby also determining the rival male’s cross-contamination level – or, we identified them by their unique colour-code, which was also used to distinguish between the males during the fight for the subset of 208/347 fights observed for male and worker behaviour (see below; note that during observation it was unknown to the observer, which colour reflected which male treatment). For colour-coding, we applied a tiny dot of metallic email colour (Revell) on the male’s dorsal abdomen (gaster) using a micro dissection needle holder with a fine tungsten needle (1 μm tip diameter), and let the colour dry for multiple hours before start of the experiment. Colour coding was used for 109 high-dose fights and all 111 low-dose fights (Table S1), As the proportion of infected individuals being the loser of a fight per treatment group did not differ between male identification method (PCR vs colour), we (i) concluded that colour-coding of the males did not alter fight outcome, and (ii) pooled the two methods for analysis of who was the winner or loser in decided fights (note that for 8/48 PCR identifications this distinction was not possible, as detailed below, Table S2, so that the identity of the healthy vs infected male winning or losing could only be resolved in 96/112 fights).

### Spore load quantification by ddPCR

In 48 fights of differently-treated males with one male being dead and the other alive, we determined whether it was the infected or the healthy male that had lost the competition. To this end, we froze both rivals after the fight, extracted DNA and ran a droplet digital PCR (ddPCR; Bio-Rad) for direct quantification of their spore load.

For DNA extraction, the samples were homogenized using a TissueLyser II (Qiagen) with a mixture of one 2.8 mm ceramic (Qiagen), five 1 mm zirconia (BioSpec Products) and ~100 mg of 425-600 μm glass beads, acid washed (Sigma-Aldrich) in 50 μl of nuclease-free water (Sigma). Homogenization was carried out in two steps (2 x 2 min at 30 Hz). After the first 2 min the tube racks were rotated to ensure uniform disruption and homogenization of all samples. In case the samples were not uniformly crushed yet, the homogenization was repeated. DNA extraction was performed using the DNeasy 96 Blood & Tissue Kit (Qiagen) following the manufacturer’s instructions with a final elution volume of 50 μl. As the used *M. robertsii* strain (ARSEF 2575) is fluorescently labelled, with each spore carrying a single plasmid with a single copy of the *mRFP1* gene (Fang *et al*. 2006), we could determine absolute spore numbers of our samples by targeting the *mRFP1* gene. Primers and probe were designed to bind to the *mRFP1* gene (gene bank accession number: KX176868.1; mRFP1_Forward: 5’-CTGTCCCCTCAGTTCCAGTA, mRFP1_Reverse: 5’-CCGTCCTCGAAGTTCATCAC, mRFP1_probe: 5’[6FAM]AGCACCCCGCCGACATCCCCG[BHQ1]) using Primer3Plus software (Untergasser *et al*., 2007) and were confirmed to only amplify the gene of interest.

The ddPCR reactions with a total volume of 22 μl were prepared as follows: 11 μl of 2x ddPCR Supermix for Probes (Bio-Rad), 900 nM of both primers (Sigma Aldrich) and 250 nM probe (Sigma Aldrich), 10 U of both enzymes EcoRI-HF and HindIII-HF (New England Biolabs), 5.27 μl nuclease-free water (Sigma) and 2.2 μl of template DNA. Droplets were generated according to manufacturer’s instructions with the QX200 Droplet Generator. The amplification program was initiated with a first step at 95 °C for 10 min to activate the DNA polymerase, followed by 40 cycles of 30 sec at 94 °C and 60 sec at 56 °C, and finished with 98 °C for 10 min to stop the reaction. For the entire program the ramp rate was set to 2 °C/sec. After amplification droplets were analysed on the QX200 Droplet Reader (Bio-Rad) for the readout of positive and negative droplets.

Data analysis was done using the QuantaSoft Analysis Pro Software (Version 1.0.596; Bio-Rad). The threshold was manually set to 3000. Overloaded samples (low amount of negative droplets) were diluted and re-run. Each run included a H2O negative control (which in 2 of 6 cases showed a single positive droplet). We therefore only considered males that had at least 2 positive droplets in the PCR to be above threshold and to have confirmed spore load (either being the exposed male or by cross-contamination from the exposed male). In 8/48 fights we were not able to differentiate between the healthy and infected male as the fungal loads of both males were below detection threshold. These fights could thus not be included in analyses, in which identity of the males was required.

### Observation of male and worker aggression

We quantified male and worker aggressive actions by behavioural scan sampling for a subset of 97 high-dose and all 111 low-dose fights (including their respective control fights of two healthy males; see Tables S1, S3: all of these fights included colour-coded males, total N = 208 observed fights). Observers were blind for both fight combination (healthy-healthy vs healthy-infected) and individual male treatment (colour code did not reveal treatment). A total of 18 scans (each taking one to several seconds) were performed per fight, one every 30 min for the first 8h and one last one after 24h, when we also determined fight outcome. For each male, we determined the number of aggressive interactions performed towards the other male, which included several behaviours (which, if occurring simultaneously, were each counted). These were *biting, holding* (grabbing the rival with the mandibles) and *besmearing* (male bending its gaster tip to apply hindgut secretion onto the rival, often combined with holding). We also counted the number of aggressive interactions that workers performed towards either of the males. Worker aggression combined *biting, carrying* (male is held with the mandibles and carried around), *dragging* (male is held and pulled over the ground), and *dismembering* (body parts are intensively attacked or already bitten off).

### Determination of male mortality risk in the absence of fighting

To determine the *baseline mortality* of the infected males (and their healthy controls) in the absence of fighting, we kept males of each infection stage and level with five of its colony’s workers, but no second male, for 24h and determined their survival. We used a total of 268 males as detailed in Table S4.

### Activation of the immune system in mild and severe infection

We determined the immune gene activation that males show in the middle and at the end of the fighting period for 57 additional colour-marked males following treatment with the sham solution, the low or the high pathogen dose, and keeping them individually for 12 or 24h after exposure (N=9-10 males per time point and dose combination, Table S6). For each male, we determined the expression levels of three different immune genes, involved in the cellular and humoral immune response against fungal infection: (i) Relish (*Rel*) – a transcription factor in the immune deficiency pathway (IMD) (Myllymäki, Valanne and Rämet, 2014; Sheehan, Farrell and Kavanagh, 2020), (ii) the shared homolog of the Melanisation Protease 1 and 2 (*MP1* and *MP2*) from *Drosophila melanogaster*, and the Prophenoloxidase-activating enzyme (*PPOAE*) from *D. mauritiana*, and which we refer to as Prophenoloxidase(PPO)-activating factor (*PPOAF*), a gene that is involved in melanisation as it activates *PPO* (Gillespie, Kanost and Trenczek, 1997; Cerenius and Söderhäll, 2004; Cerenius, Lee and Söderhäll, 2008), and (iii) the antimicrobial peptide Defensin (*Def*) that is activated by the Toll-pathway and destroys fungal cell walls (Viljakainen and Pamilo, 2005, 2008). We used the elongation factor 1-alpha (*EF1*) as housekeeping gene (Klein *et al*., 2016).

Males were individually snap-frozen in 1.5ml Safe-Lock Tubes (Eppendorf), their RNA extracted and transcribed to cDNA before performing gene expression analysis using ddPCR. Total RNA was extracted using the Maxwell^®^ RSC instrument together with the Maxwell^®^ simplyRNA Tissue Kit (Promega) according to the manufacturer’s instructions with a final elution volume of 60 μl. Prior to RNA extraction individuals were homogenized in 200 μl homogenization buffer including 4 μl 1-Thioglycerol using a TissueLyser II (Qiagen) with a mixture of five 1 mm zirconia (BioSpec Products) and ~100 mg of 425-600 μm glass beads, acid washed (Sigma-Aldrich). Homogenization was carried out in two steps (2 x 2 min at 30 Hz). After the first 2 min the tube racks were rotated to ensure uniform disruption and homogenization of all samples. In case the samples were not uniformly crushed yet, the homogenization was repeated. To ensure the complete removal of residual DNA contamination an additional DNase-I treatment (Sigma-Aldrich) step was performed prior to the reverse transcription. The cDNA synthesis was performed using the iScript cDNA synthesis kit (Bio-Rad) according to manufacturer’s instructions.

To analyse the expression patterns of the immune genes, we used multiplex ddPCR assays, each targeting one immune gene and the housekeeping gene. Target sequences were identified by blasting published annotated sequences of ant species (*Atta cephalotes* and *Temnothorax longispinosus*) against the *Cardiocondyla obscurior* genome (Errbii *et al*., 2021), accessible under GenBank number GCA_019399895.1. Primers were designed using *Primer3Plus* (Untergasser *et al*., 2007) and *Multiple Primer Analyzer* (https://www.thermofisher.com) software. We used the following primers and probe for the housekeeping gene: EF1_Forward: 5’-ATTGGAACAGTACCCGTTGG, EF1_Reverse: 5’-CACCCTTCGGTGGGTTATTT, EF1_probe: 5’[HEX]ACCTGGTATGGTCGTTACCTTTGCACCCGT[BHQ1]; the Relish gene: Rel_Forward: 5’-ACGGATTTAGGATGGACACC, Rel_Reverse: 5’- TTGGTGGCTTCCTT CAACA, Rel_probe: 5’[6FAM]TGCTCTCTTGTGCAGACTGGCGCAGA[BHQ1]; the PPOAF gene: PPOAF_Forward: 5’- TGCTGCTCACTGTATCAAGG, PPOAF_Reverse: 5’-TCTGTTTCAGTGTCGGTGTC, PPOAF_probe: 5’ [6FAM]ACTGGCGTCTGACCAGCGTCCGT[BHQ1]; and the Defensin gene: Def_Forward: 5’-ACGGGCCTACTTACGAATTG, Def_Reverse: 5’-CGCAA GCACTATGGTTGATG, Def_probe: 5’[6FAM]CGAAGAGGAGCCGTCACACC TGACGC[BHQ1].

The ddPCR reactions with a total volume of 22 μl were prepared as follows: 11 μl of 2x ddPCR Supermix for Probes (Bio-Rad), 900 nM of each primer (Sigma-Aldrich) for the immune gene of interest as well as the housekeeping gene and 250 nM of the respective probes (Sigma Aldrich), 3.74 μl nuclease-free water (Sigma) and 2.2 μl of cDNA. The cDNA was sonicated (15 sec) before being added to the reagent mix. All further steps were performed as described above, except that the amplification program was slightly modified to improve the separation between positive and negative droplets. The amplification program was initiated with a first step at 95 °C for 10 min, followed by 50 cycles of 30 sec at 94 °C and 50 cycles of 2 min at 56 °C, and finished with 98 °C for 10 min. All PCR steps were carried out with a ramp rate of 1°C/sec. The thresholds were set manually as follows: *EF1* 4000, *PPOAF* 1500, *Rel* 2000 and *Def* 4000. Immune gene expression was normalized to the reference gene *EF1*.

### Statistical analysis

All statistical analyses were performed in the program ‘R’ version 4.0.3 (R Core Team, 2020) and all reported p-values are two-sided. Unless otherwise stated, we used a generalized mixed modelling approach (GLMM) (‘lme 4’(Bates *et al*., 2015)) in which the significance of the model predictors were estimated by comparing each model to a null model only containing the intercept using Likelihood Ratio (LR) tests (Bolker *et al*., 2009). We assessed model assumptions (residual normality and heterogeneity and no overdispersion) with ‘DHARMa’ (Hartig, 2020) and checked model stability and the presence of influential data points (Cook’s distance and dfbetas). Whenever multiple inferences were made from the same dataset, we corrected the overall p-values for multiple testing with the Benjamini-Hochberg procedure to protect against a false discovery rate of 0.05 (Benjamini and Hochberg, 1995). We report two-sided, corrected p-values. All graphs were made using the ‘ggplot2’ package (Wickham *et al*., 2018).

#### Fight outcome

To determine whether the fight outcome (one male dead, both males dead, or both males alive) differed between fight combinations (healthy-healthy vs healthy-infected) for each scenario (early stage low and high dose, as well as late stage high and low dose), we ran logistic regressions with binomial error term and logit-link function for each outcome and implemented fight outcome as response variable and fight combination as predictor variable (multiple testing adjustment for 3 comparisons each; Figs. 1a,b, Table S2) and included ‘colony’ as a random effect. To further test whether early- or late-stage infected males were losing more often against the healthy rival than expected by chance, we used χ^2^-tests or, if the minimum expected frequency was less than 5, Fisher’s exact tests (Figs. 1a,b).

#### Male aggression performed towards its rival

For fights with one healthy and one infected male we summed the number of male aggressive behaviours observed against its rival and analysed them as count data. For each fight scenario (early stage low and high dose, late stage high and low dose), we ran GLMMs with negative binomial error term and log-link function as the data was over-dispersed, with male infection status (healthy vs infected) as predictor and the number of male aggressive acts as response. ‘Colony’ and ‘experimental replicate’ were included as random effects (multiple testing adjustment for 2 comparisons each). While the statistics are performed with the raw data, we visualise the infected male’s aggression in comparison to its healthy rival (by subtracting the infected male’s aggression score from that of the healthy male for each fight; Figs. 1c,d; Table S3). We ran the same models also for the control fights between the two healthy males (N=46 early and N=50 late control fights, Table S1; low and high doses pooled, as both males in the control fights had only received sham treatment across dosages), by randomly assigning the two males per fight into either male A or male B, and iterating the model 100 times. None of the models showed a significant difference between male aggression of males categorized randomly into A and B males, confirming that male aggression was also balanced in the control fights.

#### Worker aggression towards males

For fights with one healthy and one infected male we summed the number of worker aggressive events against each male (count data). To test if the infected vs healthy males received a different level of aggression by the workers, we ran GLMMs with Poisson error terms and log-link function with male infection status (healthy vs infected) as predictor and the number of worker attacks as response. ‘Colony’ and ‘experimental replicate’ were included as random effects (multiple testing adjustment for 2 comparisons each). Statistics were run on the raw data, while we visualise the difference of worker aggression towards the infected males compared to that towards the healthy male of the same fight by subtracting the aggression towards the infected male from that directed towards the healthy male (Figs. 1e,f; Table S3). Again, we ran the same models also for the 46 early and 50 late control fights, with randomized categorisation of the two males per replicate into male A or B, and 100 times iteration of running the models, and corrected for multiple testing. For the early fights, 2/100 iterations showed a significant difference between worker aggression received by male A vs B. In the late fights, 16/100 iterations showed imbalanced worker aggression, which was mostly due to one influential fight, in which workers aggressed one of the males –the later loser– 21-fold more than in the other control fights (15 attacks vs a mean ± SD of 0.69 ± 1.36 attacks per male in all remaining late control fights), with always the category A or B being more aggressed to which this highly-attacked male was randomly assigned to (in our set of iterations 41% A and 59% B). Exclusion of this special fight led to only 7/100 of the iterations showing significant imbalance of worker aggression. Similar to fights between differentially-aged males (Cremer *et al*., 2012), overproportional worker aggression against the later loser was therefore also possible in fights between the two similarly-aged healthy males in our experiment, yet without any predictable pattern between males A and B.

#### Male mortality in the absence of fighting

To determine if the baseline risk of the males to die during the 24h fight period in the absence of fighting was higher for the infected males than their respective healthy male control, we ran a logistic regression with binomial error term and logit-link function with male survival (dead vs alive) as response and male infection status (healthy vs infected) as predictor for each dose and stage of infection (Figs. 1g,h, S1; Table S4).

#### Costs of fighting

To determine the additional costs of fighting of infected males, we compared the survival of non-fighters (baseline risk) to the survival of fighting males for each dose and infection stage. We considered fights with one healthy and one infected male that either ended decided (one dead) or both males being dead and calculated the proportion in which the infected males were dying and compared it to the proportion of male mortality in the absence of fighting using χ^2^-tests or, if the minimum expected frequency was less than 5, Fisher’s exact tests (Figs. 1g,h; Table S5; multiple testing adjustment for 4 comparisons each).

#### Immune gene activation

To test whether the gene expression differed between infected and healthy males at 12h and 24h after exposure for both the low and high dose, we compared their normalised gene expression values (value of immune gene / value of housekeeping gene) by Wilcoxon-Tests for independent samples (multiple testing adjustment for 6 comparisons). For visualisation (Figs. 2, S2), we show the difference of the infected males to their respective healthy male control, by subtracting the median of the healthy males from each infected male.

## Declarations

## Acknowledgements

We are thankful to Mike Bidochka for the fungal strain, Lukas Schrader for sharing the *C. obscurior* genome data for primer development, the Lab Support Facility of ISTA for general laboratory support and help with the permit approval procedures, and the Finca El Quinto for letting us collect ants on their property. We thank the Social Immunity Team at ISTA for help with ant collection and experimental help, in particular Elina Hanhimäki and Marta Gorecka for behavioural observation, and Elisabeth Naderlinger for spore load PCRs. We further the Social Immunity Team and Jürgen Heinze for continued discussion and comments on the manuscript.

## Ethics

*C. obscurior* is an unprotected invasive species. Ant collection followed the rules for Access and Benefit-Sharing following the Nagoya protocol and was granted by the Spanish Ministry of Agriculture, Fisheries and Environment (ABSCH-IRCC-ES-237603 ESNC2), as well as the Council of Tenerife (Expte: AFF 199/17). All experimental work is in line with European law and institutional guidelines.

## Data availability

The datasets generated and analysed in this study will be made publicly available upon acceptance.

## Competing interests

The authors declare they have no competing interests.

## Funding

This project received funding from the European Research Council (ERC) under the European Union’s Horizon 2020 research and innovation programme (grant agreement No 771402 to SC). The funding body had no role in the design of the study and collection, analysis, and interpretation of data and in writing the manuscript.

## Authors’ contributions

SM and SC conceived the study. The experiments were performed by SM. JK and AVG established the molecular methods. JK collected the data for the immune gene expression. SM curated and analysed all data, with JK performing the randomization statistics. SM prepared the figures. SM and SC designed the data visualisation and wrote the manuscript. All authors read and approved the final manuscript.

## Notes

### Competing Interest Statement

The authors have declared no competing interest.

## References

Abe, J., Kamimura, Y. and Shimada, M. (2005) ‘Individual sex ratios and offspring emergence patterns in a parasitoid wasp, *Melittobia australica* (Eulophidae), with superparasitism and lethal combat among sons’, Behavioral Ecology and Sociobiology, 57(4), pp. 366–373. doi:10.1007/s00265-004-0861-y.

Alexander, J. and Stimson, W.H. (1988) ‘Sex hormones and the course of parasitic infection’, Parasitology Today, 4(7), pp. 189–193. doi:10.1016/0169-4758(88)90077-4.

Angelone, S. and Bidochka, M.J. (2018) ‘Diversity and abundance of entomopathogenic fungi at ant colonies’, Journal of Invertebrate Pathology, 156(June), pp. 73–76. doi:10.1016/j.jip.2018.07.009.

Baer, B., Krug, A., Boomsma, J.J. and Hughes, W.O.H. (2005) ‘Examination of the immune responses of males and workers of the leaf-cutting ant *Acromyrmex echinatior* and the effect of infection’, Insectes Sociaux, 52, pp. 298–303. doi:10.1007/s00040-005-0809-x.

Barribeau, S.M. and Otti, O. (2020) ‘Sexual Reproduction and Immunity’, eLS, (Box 1), pp. 1–10. doi:10.1002/9780470015902.a0028146.

Bates, D., Mächler, M., Bolker, B.M. and Walker, S.C. (2015) ‘Fitting linear mixed-effects models using lme4’, Journal of Statistical Software, 67(1). doi:10.18637/jss.v067.i01.

Benjamini, Y. and Hochberg, Y. (1995) ‘Controlling the False Discovery Rate: A Practical and Powerful Approach to Multiple Testing’, Journal of the Royal Statistical Society: Series B (Methodological), 57(1), pp. 289–300. doi:10.1111/j.2517-6161.1995.tb02031.x.

Bolker, B.M. et al. (2009) ‘Generalized linear mixed models: a practical guide for ecology and evolution’, Trends in Ecology and Evolution, 24(3), pp. 127–135. doi:10.1016/j.tree.2008.10.008.

Boomsma, J.J. (2013) ‘Beyond promiscuity: Mate-choice commitments in social breeding’, Philosophical Transactions of the Royal Society B: Biological Sciences, 368(1613). doi:10.1098/rstb.2012.0050.

Boomsma, J.J., Baer, B. and Heinze, J. (2005) ‘The evolution of male traits in social insects.’, Annual review of entomology, 50, pp. 395–420. doi:10.1146/annurev.ento.50.071803.130416.

Casillas-Pérez, B., Pull, C.D., Naiser, F., Naderlinger, E., Matas, J. and Cremer, S. (2022) ‘Early queen infection shapes developmental dynamics and induces long-term disease protection in incipient ant colonies’, Ecology Letters, 25(1), pp. 89–100. doi:10.1111/ele.13907.

Cerenius, L., Lee, B.L. and Söderhäll, K. (2008) ‘The proPO-system: pros and cons for its role in invertebrate immunity’, Trends in Immunology, 29(6), pp. 263–271. doi:10.1016/j.it.2008.02.009.

Cerenius, L. and Söderhäll, K. (2004) ‘The prophenoloxidase-activating system in invertebrates’, Immunological Reviews, 198, pp. 116–126. doi:10.1111/j.0105-2896.2004.00116.x.

Clutton-Brock, T.H. (1984) ‘Reproductive effort and terminal investment in iteroparous animals’, The American Naturalist, 123(2), pp. 212–229.

Clutton-Brock, T.H., Albon, S.D., Gibson, R.M. and Guinness, F.E. (1979) ‘The logical stag: Adaptive aspects of fighting in red deer (Cervus elaphus L.)’, Animal Behaviour, 27(PART 1), pp. 211–225. doi:10.1016/0003-3472(79)90141-6.

Cremer, S. et al. (2008) ‘The Evolution of Invasiveness in Garden Ants’, PLOS ONE, 3(12), p. e3838. doi:10.1371/journal.pone.0003838.

Cremer, S. (2019) ‘Social immunity in insects’, Current Biology, 29(11), pp. R458–R463. doi:10.1016/j.cub.2019.03.035.

Cremer, S., Armitage, S.A.O. and Schmid-Hempel, P. (2007) ‘Social immunity.’, Current Biology, 17(16), pp. R693–R702. doi:10.1016/j.cub.2007.06.008.

Cremer, S. and Heinze, J. (2003) ‘Stress grows wings: Environmental induction of winged dispersal males in Cardiocondyla ants’, Current Biology, 13(3), pp. 219–223. doi:10.1016/S0960-9822(03)00012-5.

Cremer, S., Suefuji, M., Schrempf, A. and Heinze, J. (2012) ‘The dynamics of male-male competition in Cardiocondyla obscurior ants’, BMC Ecology, 12(7), pp. 11–15. doi:10.1186/1472-6785-12-7.

Dantzer, R. (2001) ‘Cytokine-induced sickness behavior: mechanisms and implications. Ann NY Acad Sci Cytokine-Induced Sickness Behavior’, Annals of the New York Academy of Sciences, (933), pp. 222–234. doi:10.1111/j.1749-6632.2001.tb05827.x.

Eisenegger, C., Haushofer, J. and Fehr, E. (2011) ‘The role of testosterone in social interaction’, Trends in Cognitive Sciences, 15(6), pp. 263–271. doi:10.1016/j.tics.2011.04.008.

Errbii, M. et al. (2021) ‘Transposable elements and introgression introduce genetic variation in the invasive ant Cardiocondyla obscurior’, Molecular Ecology, 30(23), pp. 6211–6228. doi:10.1111/mec.16099.

Fang, W., Pei, Y. and Bidochka, M.J. (2006) ‘Transformation of *Metarhizium anisopliae* mediated by *Agrobacterium tumefaciens*’, Canadian Journal of Microbiology, 52(7), pp. 623–626. doi:10.1139/W06-014.

Gerloff, C.U., Ottmer, B.K. and Schmid-Hempel, P. (2003) ‘Effects of inbreeding on immune response and body size in a social insect, Bombus terrestris’, Functional Ecology, 17(5), pp. 582–589. doi:10.1046/j.1365-2435.2003.00769.x.

Gillespie, J.P., Kanost, M.R. and Trenczek, T. (1997) ‘Biological mediators of insect immunity’, Annual Review of Entomology, 42, pp. 611–643. doi:10.1146/annurev.ento.42.1.611.

Hajek, A.E. and St Leger, R.J. (1994) ‘Interactions between fungal pathogens and insect hosts’, Annual Review of Entomology, 39(1), pp. 293–322. doi:10.1146/annurev.ento.39.1.293.

Hamilton, W.D. and Zuk, M. (1982) ‘Heritable true fitness and bright birds: A role for parasites?’, Science, 218(4570), pp. 384–387. doi:10.1126/science.7123238.

Hart, B.L. (1988) ‘Biological basis of the behavior of sick animals’, Neuroscience and Biobehavioral Reviews, 12(2), pp. 123–137. doi:10.1016/S0149-7634(88)80004-6.

Hartig, F. (2020) ‘DHARMa: Residual Diagnostics for Hierarchical Regression Models’, The Comprehensive R Archive Network, pp. 1–26. Available at: http://florianhartig.github.io/DHARMa/%0Ahttps://cran.r-project.org/package=DHARMa.

Heinze, J. (2017) ‘Life-history evolution in ants: The case of *Cardiocondyla*’, Proceedings of the Royal Society B: Biological Sciences, 284(1850). doi:10.1098/rspb.2016.1406.

Heinze, J., Cremer, S., Eckl, N. and Schrempf, A. (2006) ‘Stealthy invaders: the biology of *Cardiocondyla* tramp ants’, Insectes Sociaux, 53(1), pp. 1–7. doi:10.1007/s00040-005-0847-4.

Heinze, J. and Hölldobler, B. (1993) ‘Fighting for a harem of queens: Physiology of reproduction in *Cardiocondyla* male ants’, Proceedings of the National Academy of Sciences of the United States of America, 90(18), pp. 8412–8414. doi:10.1073/pnas.90.18.8412.

Heinze, J., Hölldobler, B. and Yamauchi, K. (1998) ‘Male competition in *Cardiocondyla* ants’, Behavioral Ecology and Sociobiology, 42(4), pp. 239–246. doi:10.1007/s002650050435.

Helft, F., Monnin, T. and Doums, C. (2015) ‘First evidence of inclusive sexual selection in the ant *Cataglyphis cursor*: worker aggressions differentially affect male access to virgin queens’, Ethology, 121(7), pp. 641–650. doi:10.1111/eth.12376.

Hillgarth, N. and Wingfield, J.C. (1997) ‘Testosterone and Immunosuppression in Vertebrates: Implications for Parasite-Mediated Sexual Selection’, Parasites and Pathogens, pp. 143–155. doi:10.1007/978-1-4615-5983-2_7.

Hölldobler, B. and Wilson, E.O. (1990) The Ants. Harvard University Press.

Holway, D.A., Lach, L., Suarez, A. V., Tsutsui, N.D. and Case, T.J. (2002) ‘The causes and consequences of ant invasions’, Annual Review of Ecology and Systematics, 33(Redford 1987), pp. 181–233. doi:10.1146/annurev.ecolsys.33.010802.150444.

Hughes, W.O.H., Eilenberg, J. and Boomsma, J.J. (2002) ‘Trade-offs in group living: transmission and disease resistance in leaf-cutting ants’, Proceedings of the Royal Society B, (August), pp. 1811–1819. doi:10.1098/rspb.2002.2113.

Kemp, D.J. and Wiklund, C. (2001) ‘Fighting without weaponry: A review of male-male contest competition in butterflies’, Behavioral Ecology and Sociobiology, 49(6), pp. 429–442. doi:10.1007/s002650100318.

Kenis, M. et al. (2009) ‘Ecological effects of invasive alien insects’, Biological Invasions, 11(1), pp. 21–45. doi:10.1007/s10530-008-9318-y.

Kent, S., Bluthé, R.M., Kelley, K.W. and Dantzer, R. (1992) ‘Sickness behavior as a new target for drug development’, Trends in Pharmacological Sciences, 13(C), pp. 24–28. doi:10.1016/0165-6147(92)90012-U.

Kinomura, K. and Yamauchi, K. (1987) ‘Fighting and mating behaviors of dimorphic males in the ant - *Cardiocondyla wroughtoni’*, Journal of Ethology, 5(1), pp. 75–81. doi:10.1007/BF02347897.

Klein, A. et al. (2016) ‘A novel intracellular mutualistic bacterium in the invasive ant *Cardiocondyla obscurior*’, The ISME Journal, 10, pp. 376–388. doi:10.1038/ismej.2015.119.

Konrad, M. et al. (2012) ‘Social transfer of pathogenic fungus promotes active immunisation in ant colonies’, PLOS Biology, 10(4), p. e1001300. doi:10.1371/journal.pbio.1001300.

Konrad, M. et al. (2018) ‘Ants avoid superinfections by performing risk-adjusted sanitary care’, Proceedings of the National Academy of Sciences, 115(11), pp. 2782–2787. doi:10.1073/PNAS.1713501115.

Liu, L., Li, G., Sun, P., Lei, C. and Huang, Q. (2015) ‘Experimental verification and molecular basis of active immunization against fungal pathogens in termites.’, Scientific reports, 5, p. 15106. doi:10.1038/srep15106.

Liu, P., Wei, J., Tian, S. and Hao, D. (2017) ‘Male-male lethal combat in the quasi-gregarious parasitoid *Anastatus disparis* (Hymenoptera: Eupelmidae)’, Scientific Reports, (March), pp. 1–8. doi:10.1038/s41598-017-11890-x.

Loehle, C. (1995) ‘Social barriers to pathogen transmission in wild animal populations’, Ecology, 76(2), pp. 326–335.

Mazur, A. and Booth, A. (1998) ‘Testosterone and dominance in men’, Behavioral and Brain Sciences, 21(3), pp. 353–397. doi:10.1017/S0140525X98001228.

Metzler, S., Heinze, J. and Schrempf, A. (2016) ‘Mating and longevity in ant males’, Ecology and Evolution, 6(24), pp. 8903–8906. doi:10.1002/ece3.2474.

Myllymäki, H., Valanne, S. and Rämet, M. (2014) ‘The *Drosophila* Imd Signaling Pathway’, The Journal of Immunology, 192(8), pp. 3455–3462. doi:10.4049/jimmunol.1303309.

O’Donnell, S. and Beshers, S.N. (2004) ‘The role of male disease susceptibility in the evolution of haplodiploid insect societies’, Proceedings of the Royal Society B: Biological Sciences, 271(1542), pp. 979–983. doi:10.1098/rspb.2004.2685.

Pull, C.D. et al. (2018) ‘Destructive disinfection of infected brood prevents systemic disease spread in ant colonies’, eLife, 7, pp. 1–29. doi:10.7554/elife.32073.

Pull, C.D., Hughes, W.O.H. and Brown, M.J.F. (2013) ‘Tolerating an infection: An indirect benefit of co-founding queen associations in the ant *Lasius niger’*, Naturwissenschaften, 100(12), pp. 1125–1136. doi:10.1007/s00114-013-1115-5.

Reber, A., Purcell, J., Buechel, S.D., Buri, P. and Chapuisat, M. (2011) ‘The expression and impact of antifungal grooming in ants’, Journal of Evolutionary Biology, 24, pp. 954–964. doi:10.1111/j.1420-9101.2011.02230.x.

Rosengaus, R.B., Traniello, J.F. a, Lefebvre, M.L. and Carlock, D.M. (2000) ‘The social transmission of disease between adult male and female reproductives of the dampwood termite *Zootermopsis angusticollis’*, Ethology Ecology & Evolution, 12(4), pp. 419–433. doi:10.1080/08927014.2000.9522796.

Ruiz-Gonzáles, M.X. and Brown, M.J. (2006) ‘Males vs workers: testing the assumptions of the haploid susceptibility hypothesis in bumblebees’, Behavioral Ecology and Sociobiology, 60, pp. 501–509. doi:10.1007/s00265-006-0192-2.

Schnell, A.K., Smith, C.L., Hanlon, R.T. and Harcourt, R. (2015) ‘Giant Australian cuttlefish use mutual assessment to resolve male-male contests’, Animal Behaviour, 107, pp. 31–40. doi:10.1016/j.anbehav.2015.05.026.

Schrempf, A., Moser, A., Delabie, J. and Heinze, J. (2016) ‘Sperm traits differ between winged and wingless males of the ant *Cardiocondyla obscurior*’, Integrative Zoology, 11(6), pp. 427–432. doi:10.1111/1749-4877.12191.

Schwenke, R.A., Lazzaro, B.P. and Wolfner, M.F. (2016) ‘Reproduction-Immunity Trade-Offs in Insects’, Annual Review of Entomology, 61(61), pp. 239–256. doi:10.1146/annurev-ento-010715-023924.

Shakhar, K. and Shakhar, G. (2015) ‘Why Do We Feel Sick When Infected—Can Altruism Play a Role?’, PLoS Biology, 13(10), pp. 1–15. doi:10.1371/journal.pbio.1002276.

Sheehan, G., Farrell, G. and Kavanagh, K. (2020) ‘Immune priming: the secret weapon of the insect world’, Virulence, 11(1), pp. 238–246. doi:10.1080/21505594.2020.1731137.

Sheehan, G., Garvey, A., Croke, M. and Kavanagh, K. (2018) ‘Innate humoral immune defences in mammals and insects: The same, with differences?’, Virulence, 9(1), pp. 1625–1639. doi:10.1080/21505594.2018.1526531.

Sheldon, B.C. and Verhulst, S. (1996) ‘Ecological immunology: Costly parasite defences and trade-offs in evolutionary ecology’, Trends in Ecology and Evolution, 11(8), pp. 317–321. doi:10.1016/0169-5347(96)10039-2.

Stürup, M., Baer, B. and Boomsma, J.J. (2014) ‘Short independent lives and selection for maximal sperm survival make investment in immune defences unprofitable for leafcutting ant males’, Behavioral Ecology and Sociobiology, 68, pp. 947–955. doi:10.1007/s00265-014-1707-x.

Sunamura, E. et al. (2011) ‘Workers select mates for queens: a possible mechanism of gene flow restriction between supercolonies of the invasive Argentine ant’, Naturwissenschaften, 98(5), pp. 361–368. doi:10.1007/s00114-011-0778-z.

Team, R.C. (2020) ‘R: A language and environment for statistical computing. R Foundation for Statistical Computing, Vienna, Austria.’

Ugelvig, L. V, Kronauer, D.J.C., Schrempf, A., Heinze, J. and Cremer, S. (2010) ‘Rapid antipathogen response in ant societies relies on high genetic diversity’, Proceedings of the Royal Society B: Biological Sciences, 277(1695), pp. 2821–2828. doi:10.1098/rspb.2010.0644.

Untergasser, A., Nijveen, H., Rao, X. and Bisseling, T. (2007) ‘Primer3Plus, an enhanced web interface to Primer3’, Nucleic Acids Research, 35, pp. 71–74. doi:10.1093/nar/gkm306.

Vainio, L., Hakkarainen, H., Rantala, M.J. and Sorvari, J. (2004) ‘Individual variation in immune function in the ant *Formica exsecta*; effects of the nest, body size and sex’, Evolutionary Ecology, 18(1), pp. 75–84. doi:10.1023/B:EVEC.0000017726.73906.b2.

Vestergaard, S., Butt, T.M., Bresciani, J., Gillespie, A.T. and Eilenberg, J. (1999) ‘Light and Electron Microscopy Studies of the Infection of the Western Flower Thrips *Frankliniella occidentalis* (Thysanoptera: Thripidae) by the Entomopathogenic Fungus Metarhizium anisopliae’, Journal of Invertebrate Pathology, 73(1), pp. 25–33. doi:10.1006/jipa.1998.4802.

Vidal, M., Königseder, F., Giehr, J., Schrempf, A., Lucas, C. and Heinze, J. (2021) ‘Worker ants promote outbreeding by transporting young queens to alien nests’, Communications Biology, 4(1), pp. 1–8. doi:10.1038/s42003-021-02016-1.

Viljakainen, L. and Pamilo, P. (2005) ‘Identification and molecular characterization of defensin gene from the ant *Formica aquilonia*’, Insect Molecular Biology, 14(4), pp. 335–338. doi:10.1111/j.1365-2583.2005.00564.x.

Viljakainen, L. and Pamilo, P. (2008) ‘Selection on an antimicrobial peptide defensin in ants’, Journal of Molecular Evolution, 67(6), pp. 643–652. doi:10.1007/s00239-008-9173-6.

Walker, T.N. and Hughes, W.O.H. (2009) ‘Adaptive social immunity in leaf-cutting ants’, Biology Letters, 5(4), pp. 446–448. doi:10.1098/rsbl.2009.0107.

West, S.A., Murray, M.G., Machado, C.A., Griffin, A.S. and Herre, E.A. (2001) ‘Testing Hamilton’s rule with competition between relatives’, Nature, 409(6819), pp. 510–513. doi:10.1038/35054057.

Wickham, H. et al. (2018) ‘ggplot2: Create Elegant Data Visualisations Using the Grammar of Graphics’.

Yamauchi, K. and Kawase, N. (1992) ‘Pheromonal manipulation of workers by a fighting male to kill his rival males in the ant *Cardiocondyla wroughtonii*’, Naturwissenschaften, 79(6), pp. 274–276. doi:10.1007/BF01175395.

Zuk, M., Thornhill, R., Ligon, J.D. and Johnson, K. (1990) ‘Parasites and mate choice in red jungle fowl’, Integrative and Comparative Biology, 30(2), pp. 235–244. doi:10.1093/icb/30.2.235.

