## Supplemental Tables1-6 and Figures1-2 for "Trade-offs between immunity and competitive ability in fighting ant males"

**SUPPORTING INFORMATION**
**for**

### Trade-offs between immunity and competitive ability in fighting ant males

Sina Metzler[1], Jessica Kirchner[1], Anna V Grasse[1] & Sylvia Cremer[1,*]

1) ISTA (Institute of Science and Technology Austria), Am Campus 1, 3400 Klosterneuburg, Austria

this file contains
Supporting Tables S1-S6
Supporting Figures S1-S2

**Supporting Tables**

**Table S1) Number of fights per infection dose and stage.** Sample size for all fights between an a healthy male and a male exposed to the low or high pathogen dose, and their respective control fights between two healthy males. Fight outcome after 24h was determined for all fights. A subset of fights was further observed for detailed behavioural interactions between the two males and the workers to the males. Males were identified by either individual colour codes applied before fight (for all fights with behavioural observation), or after fight by spore load PCR (for the subset of decided fights with a winner and a loser; Table S2). For the latter, also cross-contamination between males was quantified.

| Infection stage | Dose | Fight combination | Number of fights | Subset observed | Male ID by colour (or PCR) |
|---|---|---|---|---|---|
| early | low | healthy-healthy | 30 | 30 | 30 |
| | | healthy-infected | 30 | 30 | 30 |
| | high | healthy-healthy | 43 | 16 | 16 |
| | | healthy-infected | 65 | 17 | 29 (20) |
| late | low | healthy-healthy | 20 | 20 | 20 |
| | | healthy-infected | 31 | 31 | 31 |
| | high | healthy-healthy | 53 | 30 | 30 |
| | | healthy-infected | 75 | 34 | 34 (28) |

**Table S2) Statistical results of fight outcome depending on infection dose and stage.** For each fight (Table S1), it was determined if both males were still alive or had died during combat, or if there was a clear winner or loser (numbers of fights per outcome given). Sample size, test statistics and p values provided for the GLMMs between the control fights of two healthy males and fights between one infected and one healthy male. All df=1. Significant and trending p values indicated in bold (pie chart inserts in Figs.1a,b). The last column shows in how many of the decided fights between an infected and a healthy male (number and % of fights) the infected male was the loser (bars in Figs. 1a,b). Value in brackets indicates the number of fights in which we were able to clearly differentiate between the males.

| Infection stage | Dose | Fight outcome: both alive | | Fight outcome: both dead | | Fight outcome: winner/loser | | |
|---|---|---|---|---|---|---|---|---|
| | | **healthy-healthy** | **healthy-infected** | **healthy-healthy** | **healthy-infected** | **healthy-healthy** | **healthy-infected** | **Infected male losing** |
| early | low | 14 | 20 | 1 | 0 | 15 | 10 | 4 |
| | | $\chi^2$=2.462, p=0.236 | | $\chi^2$=1.462, p=0.236 | | $\chi^2$=1.724, p=0.236 | | 40% |
| | high | 21 | 17 | 4 | 8 | 18 | 40 | 34 |
| | | $\chi^2$=5.994, **p=0.043** | | $\chi^2$=0.241, p=0.623 | | $\chi^2$=4.258, **p=0.059** | | 85% |
| late | low | 10 | 16 | 2 | 7 | 8 | 8 | 3 |
| | | $\chi^2$=0.013, p=0.910 | | $\chi^2$=1.411, p=0.433 | | $\chi^2$=1.125, p=0.433 | | 37.5% |
| | high | 22 | 10 | 3 | 11 | 28 | 54 (46) | 43 |
| | | $\chi^2$=13,119, **p<0.001** | | $\chi^2$=2.710, p=0.100 | | $\chi^2$=4.936, **p=0.039** | | 93.5% |

**Table S3) Statistical results of aggression of males and workers depending on male infection dose and stage.** For each fight with detailed behavioural observation (Table S1), we quantified the aggression performed by males towards each other, as well as worker aggression towards each male. Sample size, test statistics and p values for the GLMMs between male aggression performed, and worker aggression received, by the infected compared to the healthy male. All df=1. Significant p value in bold. See Figs. 1c-f.

| Infection stage | Dose | N | Male aggression | Worker aggression |
|---|---|---|---|---|
| early | low | 30 | $\chi^2$=0.459, p=0.853 | $\chi^2$=0.035, p=0.853 |
| | high | 17 | $\chi^2$=0.073, p=0.787 | $\chi^2$=1.516, p=0.436 |
| late | low | 31 | $\chi^2$=0.261, p=0.609 | $\chi^2$=2.599, p=0.214 |
| | high | 34 | $\chi^2$=0.178, p=0.673 | $\chi^2$=8.155, **p=0.009** |

**Table S4) Baseline mortality of males in the absence of fighting depending on infection dose and stage.** Sample size of males that were observed for their mortality in the absence of fighting during the 24h period immediately after exposure (early infection stage) or 48h post exposure (late infection stage), and number of males dying during this period. See Figs. S1a,b.

| | | healthy | | infected | |
|---|---|---|---|---|---|
| **Infection stage** | **Dose** | **N observed** | **N died** | **N observed** | **N died** |
| early | low | 60 | 5 | 29 | 5 |
| | high | | | 62 | 18 |
| late | low | 53 | 5 | 25 | 9 |
| | high | | | 39 | 22 |

**Table S5) Statistical results of the additional costs of fighting.** To determine the costs of fighting of infected males, we compared the survival of non-fighters (baseline risk) to the survival of fighting males for each infection stage and level using $\chi^2$-tests and Fisher's Exact test. All df=1. Significant p values in bold. See Figs. 1g,h.

| Infection stage | Dose | |
|---|---|---|
| early | low | p=0.730 |
| | high | $\chi^2$=16.120, **p<0.001** |
| late | low | $\chi^2$=0.086, p=0.769 |
| | high | $\chi^2$=7.107, **p=0.010** |

**Table S6) Statistical results of immune gene expression of males exposed to the low or high dose 12h and 24h after exposure.** Sample size, test statistics and p values for the Wilcoxon Tests comparing gene expression of each of the three measured immune genes (each normalised to the housekeeping gene *EF1*) between the infected males of the two doses to their respective healthy control males, which were measured at the same time after exposure. Significant p-values in bold. See Figs. 2, S2.

| Treatment | Time | N | Relish | *PPOAF* | Defensin |
|---|---|---|---|---|---|
| control | 12h | 9 | | | |
| | 24h | 10 | | | |
| low | 12h | 10 | W=58, p=0.473 | W=49, p=0.780 | W=50, p=0.780 |
| | 24h | 9 | W=65, p=0.135 | W=50, p=0.720 | W=68, p=0.098 |
| high | 12h | 10 | W=77, **p=0.045** | W=64, p=0.267 | W=65, p=0.267 |
| | 24h | 9 | W=80, **p=0.009** | W=75, **p=0.027** | W=80, **p=0.009** |

#### Supporting Figures

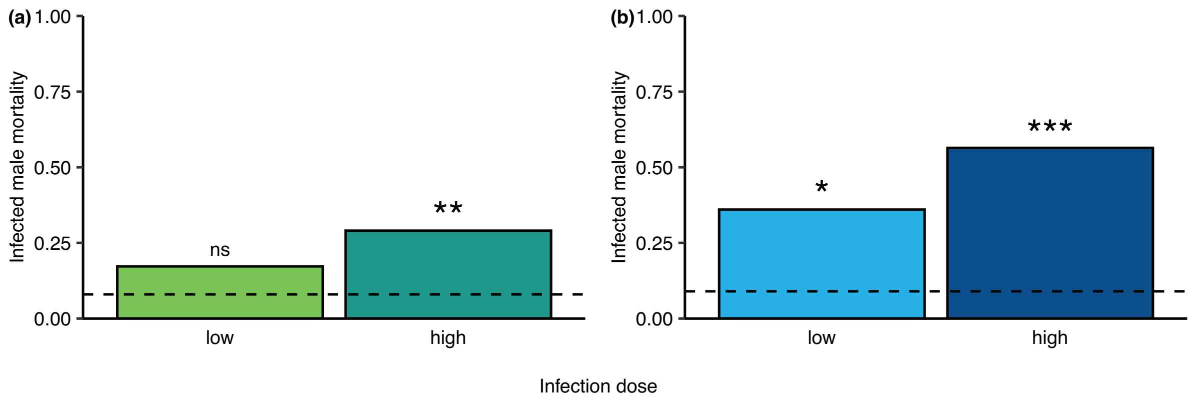


**Fig. S1) Baseline mortality risk of males in the absence of a fight in dependence of infection dose and stage.** At the early **(a;** green**)** and late **(b;** blue**)** stage of infection, the risk for an infected male to die above the risk for healthy males (dotted line) is shown for the low and high dose (darker tone reflects higher dose). Infected male mortality was significantly increased for all males except when in the early stage of infection with a low dose. Males at the late stage of a severe infection were highly moribund, not surviving the 24h in >50% even without fighting. Based on 268 males. Significant deviation to healthy male group indicated by ** p<0.01, *** p<0.001, ns=non-significant.

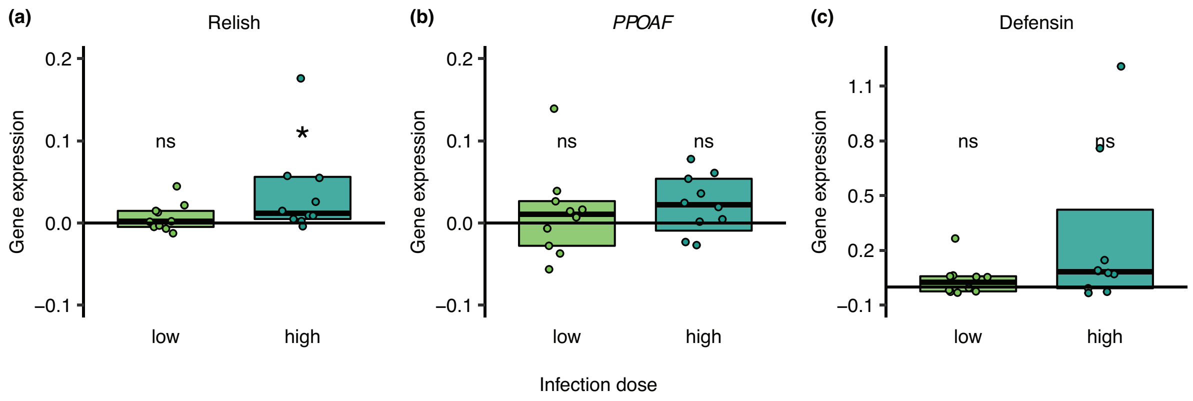


**Figure S2) Immune gene expression 12h after exposure to the low or high pathogen dose.** Expression of infected males' immune genes (**a**) Relish, (**b**) *PPOAF*, and (**c**) Defensin showed no significant activation 12h after exposure over the baseline expression of the healthy males (zero line), except for the transcription factor Relish after exposure to the high pathogen level. For each infected male we show its relative gene expression level normalised to the housekeeping gene *EF1*, relative to the median gene expression of the respective healthy control males (zero line) as individual data point, the group's median (black line) and 95% CI (box). Based on a total of 29 males (Table S6). Significant deviation to the healthy males for each dose and gene shown by * p<0.05, ns=non-significant.
